# *genomicBERT*: A Light-weight Foundation Model for Genome Analysis using Unigram Tokenization and Specialized DNA Vocabulary

**DOI:** 10.1101/2023.05.31.542682

**Authors:** Tyrone Chen, Naima Vahab, Navya Tyagi, Eleanor Cummins, Anton Y. Peleg, Sonika Tyagi

## Abstract

The genome, which serves as the inherent language directing the blueprint of life, offers significant analysis prospects by combining Natural Language Processing (NLP) and machine learning (ML). Integrating biological sequences with other digital healthcare information has potential to transform data-driven diagnostics. Large language models (LLMs) can be harnessed to decode the genomic language. This endeavor encounters three critical challenges: First, long biomolecular sequences require segmentation into smaller subunits, which is non-trivial since many biological “words” remain unknown. Second, the analysis of extended DNA sequences using LLMs demands a compute-intensive infrastructure. Third, ensuring reproducibility and reusability of modeling workflows remains an unresolved issue. To tackle these challenges, we introduce an empirical DNA tokenisation approach and a versatile, semantic-aware, genome language model —*genomicBERT*. The model is species-agnostic and operates seamlessly at the DNA or RNA levels. By introducing a reduced and specialized DNA vocabulary, our approach minimizes computational overhead and optimizes performance. Our benchmarking demonstrates that the *genomicBERT* matches or surpasses the performance of contemporary tools on the same datasets under different experimental conditions. To encourage collaboration and ease of access, we introduce *genomicBERT* as an integral component of the openly accessible conda package, *genomeNLP*. Validated across diverse case studies, *genomicBERT* lowers the barriers to decoding genomic language, relying solely on sequence data to extract meaningful insights.

**Highlights:**

- This novel model offers a compelling solution for DNA sequence analysis by significantly reducing model size and computational costs without compromising performance, setting a new standard for efficient model development.
- We demonstrate that a powerful vocabulary and tokenization method helps to derive patterns from biological sequence data while accounting for hidden semantic rules.
- Our method is agnostic to species or biomolecule type as it is data-driven. Hence, it can be applied to DNA and RNA
- We validate the important *genomicBERT* tokens by mapping back to the biologically significant motifs.
- We present a publicly available genome language modeling toolkit called *genomeNLP*, specifically designed to combine computational linguistics and genomics, enabling researchers from biology backgrounds to analyze and interpret genomic sequences effectively.

## 1 Background

Biological sequence data, encompassing nucleotide or amino acid sequences, is rich in low-level information but is commonly transformed in analysis pipelines, leading to varied degrees of information loss. In biology, such analyses can involve predicting protein structure [1], analysing images [2], identifying epigenetic features [3], and functionally annotating sequences [4]. We consider that many biological sequence data analysis pipelines irreversibly summarise data to reduce features (e.g. number of genes or variants), using the resulting subset of data to draw conclusions [5]. Conversely, this work concentrates on the primary, information-rich form of data—raw biological sequences. Multiple ways to formulate biological problems and represent biological and biomedical data in the context of machine learning exist. Biomedical images such as Magnetic Resonance Imaging (MRI) and histology data require comparatively little preprocessing, while other modalities such as gene expression data or electronic health records (EHR) often require specific preprocessing to be reformatted into an appropriate data representation [6][7]. However, the inherently sequential nature of nucleic acid and amino acid sequences makes an NLP framework intuitive.

Generally, the first step in a NLP pipeline involves segmenting a sequence into a series of smaller units for machine-compatible processing using a process called “tokenisation”. In natural languages, conventional tokenisation requires as prior knowledge the concept of a “word”. While some languages such as English can be easily split into smaller elements (or words) based on space or punctuation separation, many languages do not have this exploitable property. In addition, some languages such as Mandarin have characters which change meaning depending on the context of neighbouring characters. A custom set of grammar and user-defined rules is often used, which is specific to the language of interest as well as the context of the study [8, 9].

Similarly, in biology, sequences are commonly partitioned into user-defined blocks of *k*, referred to as *k*-mers, akin to the concept of n-grams in NLP [10]. Interestingly, the challenges in defining “words” in biological sequences align with those in character-based languages, allowing the application of the same preprocessing method to deduce “words” from the data.

The main biological tokenisation approach relies on manually defined segmentation rules, such as breaking a sequence into *k*-mers of a specified length [11] [12]. The value of *k* is typically estimated or experimented with over a range of lengths, often determined in an ad-hoc manner. Unlike human languages, where grammar and words are well-understood, biological counterparts remain elusive, making the use of *k*-mers sufficient for yielding results, even though they might not fully reflect biological reality or the context of neighboring sequences. Additionally, the computational expense of accounting for every possible combination of a *k*-mer, many of which may lack meaningful significance, presents a scalability challenge. Unlike natural languages, determining the start and end points of meaningful “words” or motifs within DNA sequences is challenging.

Compounding this problem is the known issue of out-of-vocabulary words (OOV) in NLP. In such cases, novel words or k-mers absent from the initial dataset cannot be evaluated, since conventional methods cannot reconstruct non-existing k-mers from individual characters - i.e. nucleotides or amino acids alone. Some methods attempt to mitigate the lack of biological meaning by including a range of overlapping k-mers [13] [14], but the underlying issues of defining a “biological word” as well as computational complexity remain. The segmentation problem is worsened by the length of the biological sequences. For instance, DNA sequences can be extremely lengthy, ranging to millions of base pairs (bp). Managing long DNA sequences is challenging due to repeats, long-range interactions, and sequence variants. Since DNA can be read bidirectionally, its “meaning” depends on regulatory elements (such as transcription factor binding sites or TFBS), chromatin structure, and variants. Predicting variant effects on gene expression requires capturing long-range dependencies, where distant sequence regions influence each other, sometimes across millions of base pairs, which poses a challenge for standard language models. The extensive length and features of DNA sequences result in complex data, necessitating tokenisation and modelling approaches capable of efficiently processing and interpreting such large-scale information. To obtain a quantitative data representation, the numeric representation of words is compiled after tokenisation. Frequency-based [15] or embedding approaches [16] are commonly used strategies. Tokens are represented as count values or weighted count values in frequency-based approaches. With embeddings, tokens are represented as distributed vectors in n-dimensional space, allowing users to observe the semantic relationship between “words”, which is a unique property among NLP data representations. (Table S1)

Commonly used algorithms both inside and outside language-based processing include decision tree-based models such as XGBoost [17] or Random Forest [18] and deep neural networks (DNNs) such as RNN (Recurrent Neural Network) [19] or Bidirectional Encoder Representations from Transformers (BERT) [20]. Released in 2018, the BERT language model features a bidirectional learning architecture designed to capture both syntactic and contextual information from natural language texts. BERT employs two key methods for effective learning: Next Sentence Prediction (NSP) and Masked Language Modeling (MLM). NSP helps the model to learn the likelihood of one sentence following another, while MLM involves predicting masked tokens based on their surrounding context. The model’s attention mechanism and bidirectional processing allows it to capture contextual information and dependencies between words, enhancing its semantic understanding. BERT’s implementation consists of two main stages: pretraining and fine-tuning. During pretraining, BERT is trained on large amounts of unlabeled data, which can be domain-specific. Fine-tuning then adapts the pretrained model to specific tasks, such as classification, using labeled datasets. We have considered BERT approach for modelling genomic analysis.

Previously, BERT has been implemented for DNA sequence modelling in [14] and [21]. However, their approach did use traditional k-mer based segmentation and hence, would inherit previously discussed limitations of a rule-based segmentation approach. The improvement in the form of DNABERT-2 [22] addresses the challenges of k-mer tokenisation by using Byte Pair Encoding (BPE) tokenisation [23] and used a vocabulary of size 4096. Vocabulary size will be an indication of maximum genomic sequence tokens that can be modelled. Similarly, another DNA lnaguage model called GenaLM [24] uses BPE tokeniser along with sparse attention with a vocabulary size of 32k and should be able to process up to 36k long sequences. While DNABERT-2 and GenaLM are based on BERT architecture, in HyenaDNA [25] the authors have used implicit convolutions to consider the DNA sequences at single nucleotide resolution and claimed to analyse sequences of up to 1M nucleotides.

We identified main gaps in existing language models applied to biological sequence data. First, the above mentioned BERT models are very large in size requiring millions of parameters to optimise, which can make the computing cost formidable in standards academic settings. Secondly, newer tokenisation approaches can be investigated to build high quality vocabulary of manageable size that can be customised to a given biomolecular sequence type. Additionally, existing empirical tokenisation methods have primarily focused on achieving tokens of max length up to 32 [22] and 64 [24] for DNA data but have not explored optimal lengths for DNA language models, leaving this area unaddressed in the literature. Thirdly, foundation models utilizing transformer architectures present a significant barrier to researchers from biological backgrounds due to their inherent complexity and lack of transparency. A critical need exists for a simple, accessible and automated pipeline that enables seamless file processing, model training, hyperparameter tuning, and interpretability.

Our research aims to fill these gaps by investigating the most effective token lengths, implementing a more flexible, robust and specialized vocabulary for DNA language, and optmising language model architecture to strike a trade-off between computing cost and accuracy. Our tokenisation strategy is specifically a novel approach not previously attempted in the context of the aforementioned methods. This unique DNA vocabulary and tokenization method simplifies the construction of foundation models specifically tailored for DNA sequences and can be adapted to analyze RNA levels. Furthermore, we present a user-friendly pipeline that empowers researchers from both biology and bioinformatics to conduct dynamic experimentation by systematically testing different parameter, tokenization and modeling combinations.

## 2 Results

### Data-driven Tokenisation

The tokenisation is built on the Sentencepiece and the Unigram tokeniser [27] which is a language model that considers the tokens as independent and uses a loss function to optimize the best tokens generated from a vocabulary by computing and comparing its probabilities in the corpus (See Methods). Our tokeniser has 4096 tokens in total with max token length as 16. As shown in the figure 1 (a), most of the tokens generated have a length between 5 to 9. While converting each input sequence into tokens during training of the model, a sequence of length ‘n’ can be split into average number tokens of ‘n/5’. This can be manged by the *max seq length* parameter of the model.

**Fig. 1:**
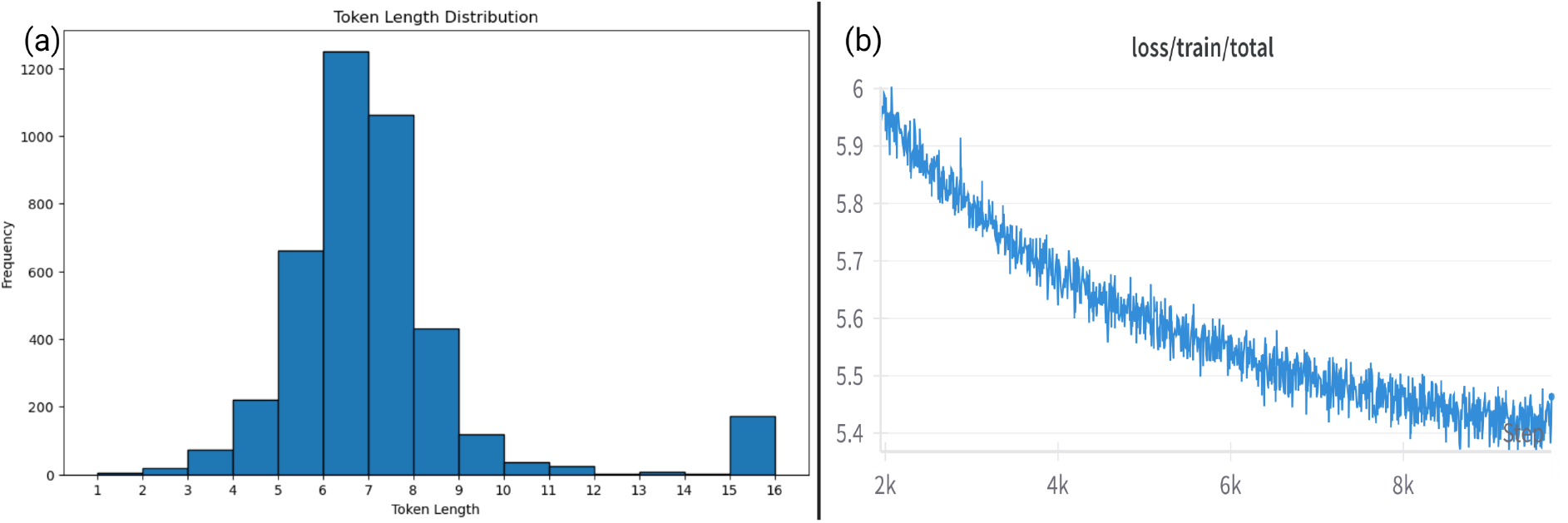
(a)Unigram tokens length analysis (b) Pretraining loss function of *genomicBERT*

### Pretraining : Development of the *genomicBERT* model

*genomicBERT* is built upon the BERT architecture and finalised with 89.2 million parameters. It follows the MosaicBERT design to take advantage of ALiBi, FlashAttention, Gated Linear Units and low precision layernorm [28]. Although the model is initially trained with a *max-seq-length* of 256 (equivalent to 1400 nucleotides), our design permits further fine-tuning on data sequences longer than 1400 nucleotides. During pretraining, the model was trained for 10k steps using 4 NVIDIA A10G GPUs of 48 vCPUs / 192 GiB Memory for 15 hours. Although the model was trained with fewer steps, our new unigram tokeniser [27] approach combined with MosaicBERT architecture excels on a variety of prediction tasks. The figure 1(b) shows the pretraining loss of the *genomicBERT*.

### Benchmarking on DNA classification as Downstream tasks

The *genomicBERT* was further fine-tuned on different classification tasks on five datasets namely, Human TFBS and lncRNA data, *E*.*coli* promoters, mouse ChIP-seq and fruit fly transcript data.

To compare efficiency of our tokeniser and model with existing DNA language models, we finetuned GenaLM [24] and DNABERT-2 [22] which are based on BPE tokenisation, on the above mentioned classification tasks. We evaluated their performance on the same datasets using training and testing ratio as 9:1. The hyperparameters for finetuning were selected for each model using a grid search approach. We report accuracy, F1 and Matthew Correlation Coefficient (MCC) metrics for all downstream tasks.

Table 1 compares the architecture of *genomicBERT* with GenaLM, and DNABERT-2 whereas, Table 2 presents the results of these three DNA foundation models on downstream classification tasks. As observed, although *genomicBERT* has significantly fewer training steps and a smaller model size, it achieves equal or higher accuracy compared to GenaLM (bert-base) and DNABERT-2 across all classification tasks. Furthermore, *genomicBERT* is able to achieve this performance with much less compute time (see num steps in Table 1) and smaller token size suggesting high quality vocabulary. Additionally, *genomicBERT* is able to perform well on cross-species data and the performance is similar to that of DNABERT-2.

**Table 1:** Comparison between different DNA Foundation models.

| Feature | *genomicBERT* | GenaLM(bert-base) | DNABERT2 |
| --- | --- | --- | --- |
| Model Size | 89.2M | 110M | 117M |
| Max sequence length | 256 | 512 | 128 |
| tokeniser approach | Unigram | BPE | BPE |
| Mask % | 10 | 15 | 15 |
| Vocab size | 4k | 32k | 4k |
| Architecture | MosaicBERT | BERT with Sparse attention | MosaicBERT |
| Hardware | 4 NVIDIA A10G GPUs | 8 or 16 NVIDIA A100 GPUs | 8 Nvidia RTX 2080Ti GPUs |
| Number of Steps | 10k | 1-2 million steps | 500k |

**Table 2:** Evaluation of Finetuning between different DNA Foundation models.

| Downstream Task | *genomicBERT* |  |  | GenaLM |  |  | DNABERT2 |  |  |
| --- | --- | --- | --- | --- | --- | --- | --- | --- | --- |
|  | Accuracy | F1 | MCC | Accuracy | F1 | MCC | Accuracy | F1 | MCC |
| Human TFBS and mRNA Identification | 0.98 | 0.98 | 0.96 | 0.51 | 0.39 | 0.06 | 0.99 | 0.99 | 0.99 |
| Human lncRNA and mRNA Identification | 0.87 | 0.87 | 0.49 | 0.75 | 0.32 | 0 | 0.95 | 0.95 | 0.91 |
| Mouse TFBS Identification | 0.96 | 0.96 | 0.90 | 0.77 | 0.67 | 0.0 | 0.99 | 0.99 | 0.97 |
| Fruit fly transcripts classification | 0.99 | 0.99 | 0.97 | 0.36 | 0.19 | 0 | 0.99 | 0.99 | 0.97 |
| Bacterial Promoter data classification | 0.79 | 0.79 | 0.48 | 0.59 | 0.34 | 0.034 | 0.79 | 0.79 | 0.58 |

### Versatile application of genomeNLP pipeline on RNA sequence data and demonstration of interpretability for biological annotation

To demonstrate the versatility of our automated genomeNLP pipeline, we applied our new tokenisation and *genomicBERT* approach to RNA fold classification problem. The positive class for this work contains pre-cursor microRNA sequences, known as pre-miRNA. Pre-miRNA sequence lengths vary but, in general, range between 50 and 300 nucleotides long [29]. Self-complementary regions bind to each other to form a hairpin like structure [Figure 2 (a)]. Pre-miRNA harbour the active form known as mature miRNA in one of its stem-loop arms which targets the genes and silences its function. Approximately 450,000 pre-miRNA sequences were downloaded in FASTA format from RNA Central [30], and while no species filtering was performed, 99% of the sequences fell within the range of 52 – 275 nucleotides, so outliers beyond this range were removed. The negative class (∼ 450,000 sequences, also sourced from RNA Central) is comprised of other small non-coding RNA sequences – a combination of snRNA (small nuclear RNAs), snoRNA (small nucleolar RNAs) and siRNA (small interfering RNA). Likewise, no species filtering was performed, but dataset was filtered on length to match the range of the positive class.

**Fig. 2:**
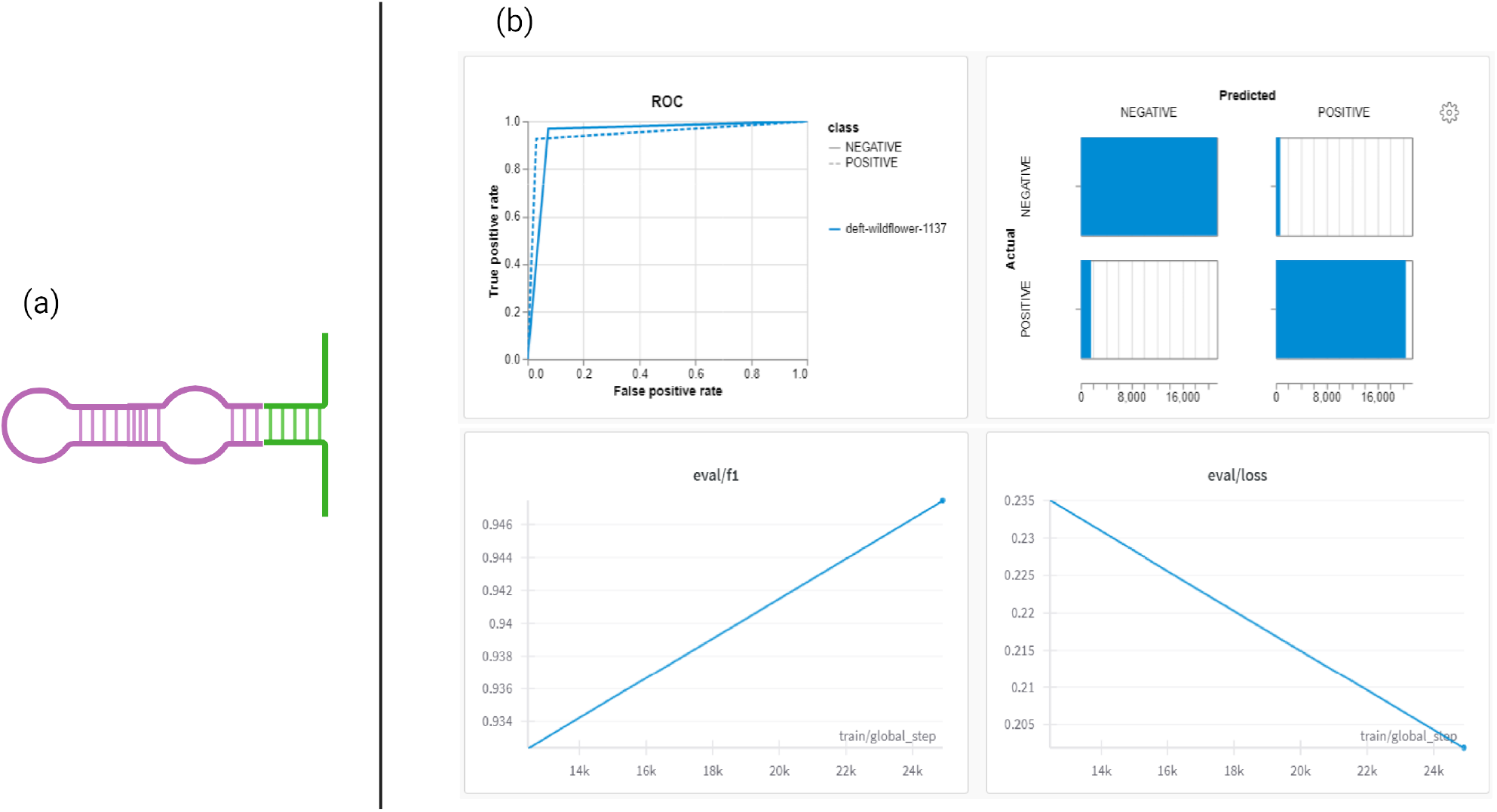
(a) Structures of miRNA with matured sequences highlighted in green; (b) Performance of *genomicBERT* on complete set of RNA data with graphs showing ROC curve, Confusion matrix, F1 curve, Loss curve respectively, from top left to right.

The genomeNLP pipeline allows multiple sequence tokenisation, token representation, and machine learning model options to run different combinations S3. For completeness we applied a more traditional kmer tokenisation and our new empirical tokeniser to this data. The best k-mer length was found to be 9 bp in the rule-based tokenisation (data not shown). On average kmerisation (k=9) of the same dataset generates approximately 10x the number of tokens as the empricial method. Kmerisation forces the creation of all possible tokens as previously described in Section 1, on the *chance* that they might be important to the model. Empirical tokenisation has already “whittled” down that set of tokens to commonly occurring tokens that have the most impact on sequence rendering, which cuts down on the inefficiency of training on features that are potentially extraneous noise. (Table S2)

The genomeNLP pipeline performed the classification and its interpretability function was used to display important tokens projected onto the sequences as a way of explaining which tokens were instrumental in deciding the classification attribution [Figure 3 (a)]. The vibrancy of the text highlight signifies the important weights attributed to that token. In the example shown here, we have added the ability to emphasize other sequences of interest. Here, the mature sub-sequences are in bold and underline. It is clear that the tokens of importance (either for or against attribution) appear densely across the miRNA sequences. Casting a qualitative eye across the examples, it also appears that the darkly highlighted sequences (i.e. those deemed of highest impact) align frequently with the mature sequences. This implies that our architecture is able to identify the most significant tokens both in biological and computational perspectives. The performance of the model is visualized in [Figure 2 (b)]

**Fig. 3:**
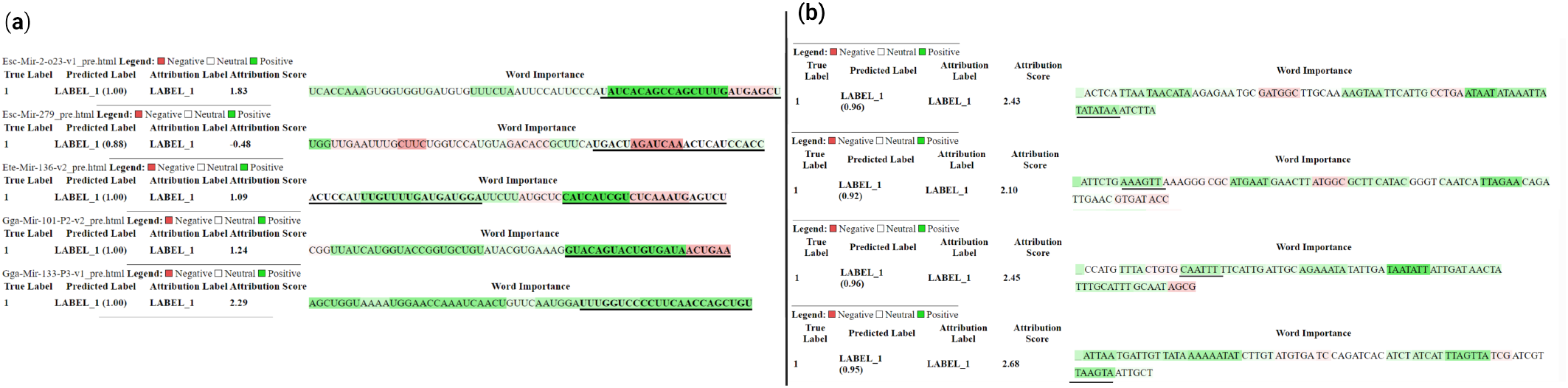
(a) Important Empirical tokens projected onto verified miRNA sequences with mature sequences underlined and bold (b) Important Empirical tokens projected onto promoter sequences. Known regulatory motifs are underlined and bold.

**Fig. 4:**
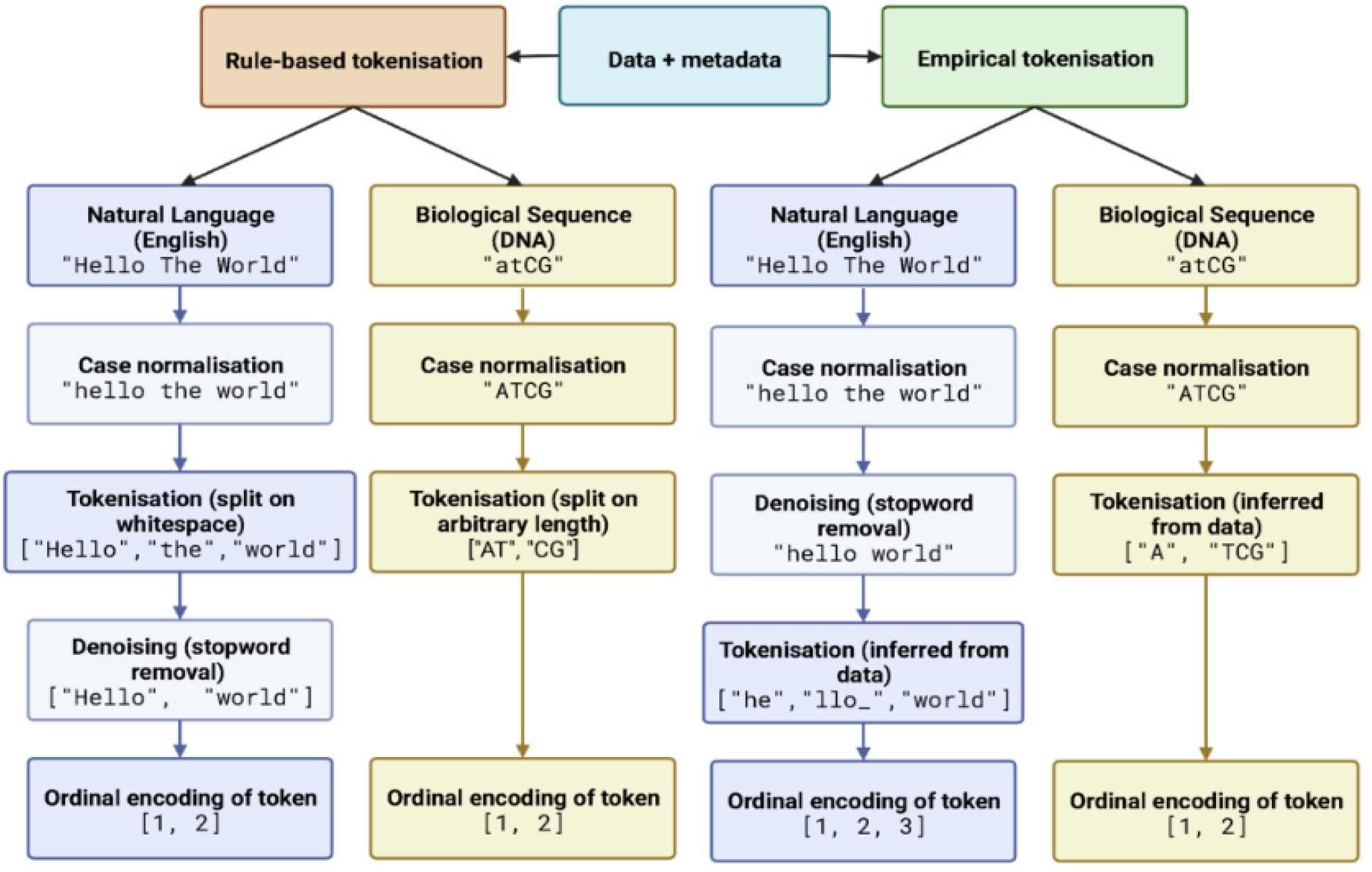
Difference between keyword based and data-driven tokenisation. Both approaches are implemented in our genomeNLP package.

Similarly, other sequence data classification tasks resulted in important tokens overlapping with serveral known motifs. For ease of visualization we show the bacterial promoter data here. Knwon regulatory motifs such as AAAGTT (sigma-28 RNA polymerase [31]), TATA boxes [32], CAATTT (Pribnow box [33]) and TAAGTA (Nkx3 [34]) are highlighted [Figure 3 (b)].

## 3 Discussion

Our DNA foundation model employs an efficient sequence tokenization approach that flexibly handles variable biological sequence lengths while preserving biological context. The enhanced BERT architecture effectively captures bidirectional and long-range interactions within genomic sequences, essential for accurate inference of hidden biological grammar. The results show that the Unigram tokenizer, when combined with the MosaicBERT architecture, performs effectively on various DNA downstream tasks, while optimising the computation needs. Furthermore, we observe that longer tokens are not necessary to represent DNA sequences. The *genomicBERT* tokenizer, which uses a maximum token length of 16, achieves performance comparable to other tokenizers, such as DNABERT2 tokeniser, and produces strong metrics across different species datasets, even without separate pretraining on these specific datasets. Therefore, our work asserts that DNA can be represented using fewer, more meaningful tokens, and the corresponding foundation models perform well across different downstream tasks and species. The superior performance of our tokenization approach stems from the probabilistic model of Unigram [27], which provides greater flexibility in generating biologically meaningful tokens at both the sub-word and character levels. This flexibility makes Unigram tokenization particularly advantageous due to its simplicity, predictability, and effectiveness in contexts involving well-defined vocabularies or character-level processing.

The effective tokenization approach in our pipeline allows us to significantly reduce model size, training time, and fine-tuning time, especially compared to state-of-the-art models. This is a crucial advancement for handling biological sequences, an area that has received limited exploration regarding the optimal number of tokens for representing biological data. Our work opens new opportunities to further investigate token efficiency while enhancing model performance with larger pretraining datasets.

Additionally, by identifying tokens with strong attribution scores for the classification task, we could pinpoint biologically significant elements within the sequences. For example, known regulatory motifs demonstrated a significant influence on the classification of promoter data. Similarly, important tokens frequently overlapped with mature miRNA sequences. Potentially novel motifs may also be annotated with this approach in an alignment-free manner [35].

Furthermore, *genomeNLP* pipeline is a user-friendly, open-source tool designed for data scientists working with biological sequence data. Available through Anaconda and Git repositories,the package supports tokenization, data transformation, and model training. Additionally, this pipeline includes visualizations that support the biological annotation of meaningful tokens and interactive exploration of training and validation metrics. This offers an intuitive means for users to explore and interpret how sequences are tokenized, with a focus on the underlying biological context.

Future work may involve extending the model to handle longer sequences more efficiently through increased computational resources and pretraining on diverse species datasets to broaden its applicability as a foundation model. The flexibility of tokenization approach naturally extends to RNA (example included) and sequence data which we aim to investigate in detail in our future work.

## 4 Conclusion

The analysis of large-scale genomic, transcriptomic, and proteomic data has long posed significant challenges due to its inherent complexity. With the exponential growth in biological datasets, there is an urgent need for modern computational methods capable of efficiently processing and extracting insights from such data. To address the computational resource constraints often encountered in research settings, we propose a novel model for DNA sequence analysis that achieves a balance between computational efficiency and predictive accuracy. This model, distinguished by its compact size, reduced training steps, and high-quality vocabulary, demonstrates performance that matches or surpasses state-of-the-art models across various downstream tasks.

Additionally, we introduce a command-line toolkit designed to streamline the application of machine learning techniques to biological sequence data. This pipeline facilitates data pre-processing and enables the identification of biologically relevant patterns in diverse sequence datasets, thereby making advanced computational methods accessible to researchers from non-computational backgrounds. To support transparency and usability, we provide metrics as interactive visualizations, enabling users to explore and evaluate model performance effectively. Detailed explanation of the methods and usage is also included as a part of the toolkit. The toolkit is designed with extensibility in mind, and we actively encourage community contributions to further enhance its capabilities. By lowering the barriers to utilizing machine learning for biological data analysis, we aim to empower researchers to pose and address novel questions within the context of large biological datasets.

## 5 Methods

The methods section is divided into two parts. First, we discuss the development of a generic foundation model for genomic sequences that we named “*genomicBERT* “. Next, we discuss our genome language toolkit called “genomeNLP” that packages *genomicBERT* along with other required steps of a NLP and deep learning based pipeline for genomic data classification.

### Data

For building the foundation *genomicBERT* model the raw FASTA files were downloaded from https://ftp.ensembl.org/pub/release-76/gtf/homo_sapiens/Homo_sapiens.GRCh38.76.gtf.gz and sequences from 24 different chromosomes were used as input for building the foundation model. The data used for validation is listed in the table(3) below.

**Table 3:** Validation data used for downstream tasks.

| Data | Description | Min, Max sequence Length in bps | Number of Samples |
| --- | --- | --- | --- |
| Regulatory sequences containing TFBS [36] | Identifies human sequences as having TFBS (non-coding regions) | 200, 1000 | 10000 |
| Long non-coding RNA and mRNA [37] | Classifies RNAs into lncRNA or mRNA sequences | 202, 9996 | 1174 |
| Mouse ChIP-seq [38] | Classifies sequences capture by ChIP sequencing | 200, 9421 | 8350 |
| Transcriptomics Data [37] | Classifies between coding and non-coding transcripts of fruit fly | 200, 9999 | 3093 |
| Promoter-non Promoter [39] | Classifies bacterial sequences as promoter and non-promoter | 81,81 | 6764 |

#### 1. Regulatory Elements Data

We focused on human regulatory sequences containing TFBS that could come from promoter or enhancer regions. The sequence locations were identified from JASPAR database (https://frigg.uio.no/JASPAR/JASPAR_TFBSs) [36] to prepare the positive set. Specifically, we extracted 100 bp region on either sides of TFBS start and end location to prepare the positive sequence data and an equal number of human coding sequence were sampled from human genome build GRCh38.

#### 2. Long non-coding Transcripts Data

We used positive lncRNA sequence data from a previously published curated set [37] where annotations from multiple lncRNA repositories were combined. Our negative data included equal number of coding transcripts from human release v34 (GRCh38.p13).

#### 3. Mouse ChIP-seq Data

We chose a publicly available mouse ChIP-seq data to investigate include data on long range regulatory interaction specifically focusing on DNase I hypersensitivity, transcription factor binding, and chromatin modifications. We downloaded this data from http://genome.ucsc.edu/cgi-bin/hgFileUi?db=mm9&g=wgEncodeSydhTfbs (GEO ID GSM912922, accessed in December 2024). We only used narrow peak data primarily focusing on long-range chromatin modifications and TFBS as positive samples and non-overlapping sequences of similar length as negative sequence class. [38]

#### 4. Fruit fly trasnscripts data

Similar to the lncRNA dataset, we curated non-coding RNAs as positive samples and coding transcripts as negative samples for fruit fly from the Ensembl database.

#### 5. Promoter Data

Promoters are important modulators of gene expression, and contain short information rich motifs that provide binding sites for regulatory proteins and co-factors. Bacterial promoter dataset was selected from a previous study [39]. This data was originally collected from RegulonDB [40] for *E. coli*. The sequences were divided into 81-bp fragments. Using CD-HIT with a threshold of 0.85, redundant sequences were removed. The dataset consists of 3382 promoters and 3382 non-promoters sequences. Our aim is to classify sequences into promoter and non-promoter sequence categories without prior knowledge of motifs.

### Preprocessing and tokenisation

There are two types of tokenisation approaches in language models; rule based and data driven. Rule based approach consists of k-mer based methods where each sentence is split into tokens of a predefined k-length with or without overlapping. The main issue with k-mer tokenisation is that it does not capture any semantic relationship between the generated tokens as well as generates much larger sized vocabulary that adds to the computational complexity of the pipeline. The data-driven tokenisation approaches consists of techniques such as BPE [41], Unigram [42] and Wordpiece [43]

With an aim to capture the most frequent and sufficiently diverse subwords, we employed the Unigram language model [27] that creates the vocabulary using a probabilistic and loss computation method. The unigram language model provides a subword segmentation that can be seen as a probabilistic mixture of characters, subwords and word segmentation [42]. BPE and Unigram are expected to have similar results, however since the unigram tokeniser is based on probabilistic model, it’s more flexible and can output multiple subword segments at character-level along with their probabilities.

The probability of a subword sequence

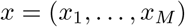

is derived as the product of the subword occurrence probabilities *p*(*x*_*i*_) using the equation (1) where V is the vocabulary set.

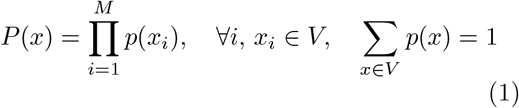

If the vocabulary *V* is predefined, the occurrence probabilities of subwords, *p*(*x*_*i*_), are estimated using the Expectation-Maximization (EM) algorithm. This approach maximizes the following marginal likelihood *L* (Formula 2), under the assumption that *p*(*x*_*i*_) are latent variables:

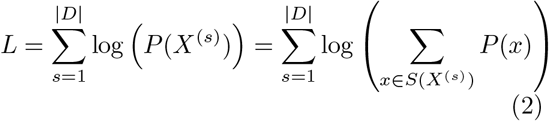

where ‘L’ is the maximum likelihood, and p(x) is the hidden variables. Figure 5(a) depicts the unigram approach for DNA sequence.

**Fig. 5:**
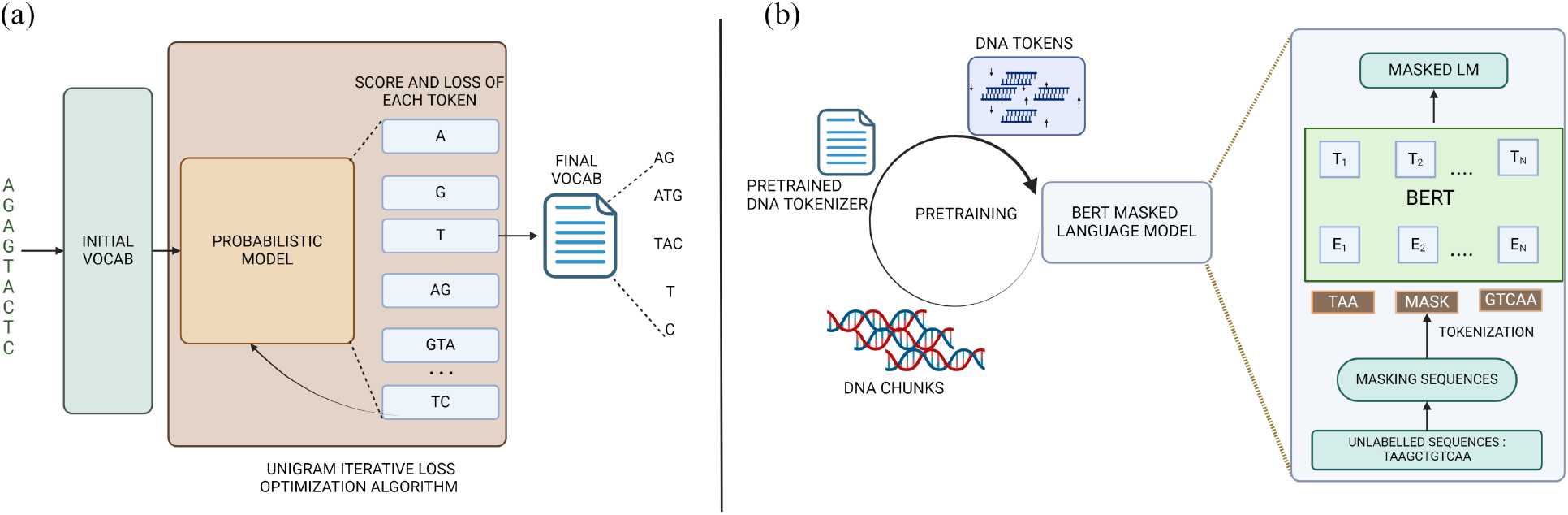
(a) Unigram tokenisation process using probabilistic model to create final vocabulary (b) *genomicBERT* architecture involving pretraining and model building.

Before tokenisation the sequences are split into chunks of 1400 nucleotides to achieve *max-seq-length* of 256 using the *genomicBERT* tokeniser. The tokeniser contains special tokens of “< *s* > “, “< */s* > “, “< *unk* > “, “< *pad* > “, “< *mask* > “. The reverse complement of the sequences are also added to the training dataset to account for any pairing of sequences. We keep pre-processing to minimal to keep the maximum biological information intact.

To construct a genome-specific tokenizer, the process begins with a single chromosome or contig. The tokenizer’s vocabulary is then iteratively refined and expanded through further pretraining on the remaining chromosomes or contigs. Specifically, we start by initialising the unigram tokeniser using sentence-piece package [26] for one of the chromosome files and generated the tokeniser and saved this as *intial vocab*. For the remaining chromosome, we initialise the unigram vocab from *intial vocab* and repeated the process for entire DNA data for dynamically updating the vocabulary. The maximum token length is set as 16 and *vocab size* as 4096. The Unigram tokenisation involves treating each word or character as a separate token. This enables character level modelling which is specially useful for analysing nucleotide level pairing interactions of sequences. Unigram is also less computationally expensive algorithm as compared to BPE and is known to produce more stable and consistent vocabulary due to the nature of the algorithm.

### Pretraining the *genomicBERT*

We use the BERT architecture as a baseline to our pipeline. MosaicBERT [28] is an enhanced version of BERT architecture to make the pretraining process faster. It implies ALiBi, FlashAttention, GatedLinearUints and Low Precision LayerNorm techniques to improve the pretraining of MLMs. The methods like rotatory positional embeddings in BERT, exhibit poor quality when applied to real data which is longer than the training data. ALiBi handles this issue by adding positional information to the attention using non learned biases which is a static set. Using this property, genomicBERT can be finetuned on different DNA sequences of length greater than 1400 nucleotides on which the model was initially pretrained. Flash attention helps to calculate the attentions in the BERT architecture in a more memory and time efficient way which help to significantly reduce the pretraining of the model. Additionally, LayerNorm operation is bfloat16 precision instead of the typical float32 precision in the MosiacBERT architecture. Due to these modifications in architecture, MosaicBERT-Base reaches higher accuracy faster than BERT-Base despite having more parameters than BERT.

The BERT MLM process involves replacing random words in a text with a [MASK] token and then learns to predict the original words. Using MosaicBERT, we have pretrained the MLM with 10% masking using the unigram tokeniser file 5(b). The detailed setting of the experiment is provided in the below Table 4. Additional details are also available in Supplementary (Figures S1 to S2)

**Table 4:** Hyper parameters used for *genomicBERT* pre-training.

| Parameter | Value |
| --- | --- |
| Mask ratio | 10% |
| Batch size | 2000 |
| Seq length | 256(1400 Ns) |
| Steps | 10k |
| $\beta_1$ | 0.9 |
| $\beta_2$ | 0.98 |
| $\epsilon$ | 1.00E-06 |
| weight decay | R1.00E-02 |
| optimizer | adamw |
| Learning rate | 5.0e-4 |
| warmup | R0.06dur |

### The genomeNLP pipeline

We packaged a pipeline with a specific focus on user-friendliness, high-level abstraction, intuitive visualisations [44] and open-source licensing. In addition, we present a systematic review of multiple case studies using our package. This package, currently available through anaconda [45] and Git repositories, is intended to be extended and built on, with any open source software developers welcome to contribute and add methods to the library. This section details about the genomeNLP pipeline is useful for data scientists working in biological data.

The pipeline employs the empirical tokenisation approach as described in the Methods (5) section. In all cases, resulting tokens were mapped to unique integers (InputIDs) as a base step before further data transformations into the final input format which the model accepts for training. Data is then assigned into training, testing, and validation buckets for future use and stored on disk as csv, json and parquet files. Each file type contains the same information, but are formatted in both human readable and machine readable formats for user convenience. The pipeline also packages simple and easy to use command line utilities for training, hyper parameter tuning and interpretation of the model. The pipeline allows one to select between multiple rule-based and deep learning models to run and compare.

For completeness we also allow for a rule-based conventional tokenisation option as part of our workflow. Since “word” delimiters are not consistently defined in biology, we arbitrarily split the sequence into equal length blocks known as k-mers on a sliding window.

### Model Interpretability

To understand how our model captures biologically meaningful information, we employed *transformers interpret* [46] to analyze the attributions of individual tokens in downstream tasks. By mapping tokens with high attribution scores to biologically significant motifs, we assessed whether the model effectively identifies key features from the dataset. Transformers Interpret [46], a package built on Captum, focuses on natural language processing (NLP) tasks, providing streamlined word attributions and visualizations for sequence classification models. An important feature of Captum is its support for Integrated Gradients (IG) [47] and Layer Integrated Gradients (LIG). LIG computes attributions by integrating gradients with respect to layer inputs or outputs along a straight-line path from baseline to the input layer activations, offering a detailed understanding of model behavior. With its user-friendly design philosophy, similar to Hugging Face’s Transformers, *transformers interpret* enables researchers to generate interpretability outputs with minimal code.

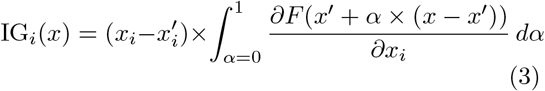

Above formula denotes IG along the i - th dimension of input X. Alpha is the scaling coefficient.

## Supporting information

Wandb

## 6 Abbreviations

BPE: Byte Pair Encoding
NLP: Natural language Processing
DL: Deep learning
BERT: Bidirectional encoder representations from transformers

## Declarations

### Ethics approval

Not applicable.

### Clinical trial number

Not applicable

### Availability of data and materials

Source DNA data was obtained from associated genomic assemblies for *Homo sapiens* **Homo sapiens.GRCh38.76**. Code associated with this project is located in https://gitlab.com/tyagilab/genomenlp. For full details on data and code please refer to Methods.

## Code availability

Git repository link and documentation: https://gitlab.com/tyagilab/genomenlp. Detailed documentation is available in the same repository and is also hosted online with case studies: https://genomenlp.readthedocs.io/en/latest/index.html. A python package is available for download and install via conda or mamba through: **mamba install -c tyronechen -c conda-forge genomenlp**.

## Authors’ contributions

T. C., N. V.: Methodology, Software, Data Curation, Writing-Original draft preparation, Writing-Reviewing and Editing, Visualisation, Investigation, Validation. N. T., E.C: Software, Writing-Original draft preparation. A. P.: Supervision. S. T.: Conceptualisation, Methodology, Writing-Original draft preparation, Writing-Reviewing and Editing, Supervision.

## Acknowledgments

This work was supported by the MASSIVE HPC facility (www.massive.org.au) and RMIT RACE Platform. We acknowledge the helpful discussions and compute resources from the Monash eResearch Platform and Monash Bioinformatics Platform. We thank Yashpal Ramakrishnaiah for helpful suggestions on package management, code architecture and documentation hosting online. We thank Jane Hawkey for advice on recovering deprecated bacterial protein identifier mappings in NCBI. We thank Andrew Perry and Richard Lupat for helping resolve an issue with the package building process. BioRender was used to create many figures in this publication. We acknowledge and pay respects to the Elders and Traditional Owners of the land on which our four Australian campuses stand.

## 7 Supplementary Material

The advantages and limitations of different tokenisation apporaches are discussed in table 1.

The pretraining metrics of *genomicBERT* are captured in the figure S1. As expected the language perplexity, cross entropy are decreasing during the pretraining. Total time taken for the pretraining is 15 hours (Figure S1 (c)). Additionally, the GPU usage for pretraining are visualized in Figure S2 (a), (b).

Furthermore, the GenomeNLP workflow is depicted in figure S3. It implies the use of a tokenisation approach, creating the dataset similar to hugging face format and finetuning using BERT with hyper parameter sweeps to select the best parameters. Final step is the evaluation and visualization using Wandb.

We also also include more analysis from *genomicBERT* finetuning for RNA data. Table S2 compares the number of tokens generated for different sets of training data from k-mer and unigram tokeniser. Figure S4 shows distribution of empirical token length in different sets of data as denoted by the S2.

**Fig. S1:**
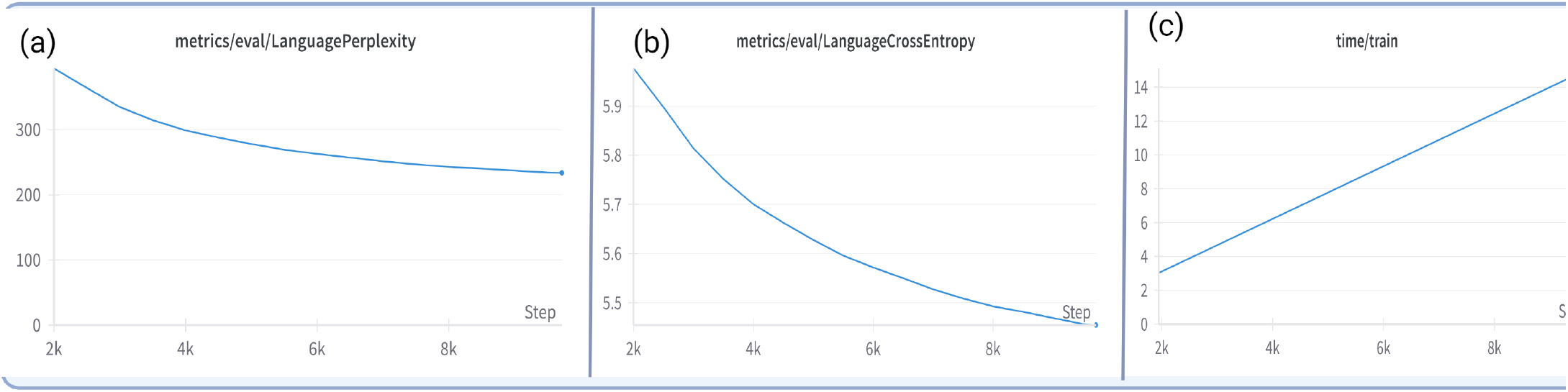
Pretraining metrics analysis.(a) Language perplexity (b) Language cross entropy (c) time taken for the analysis

**Fig. S2:**
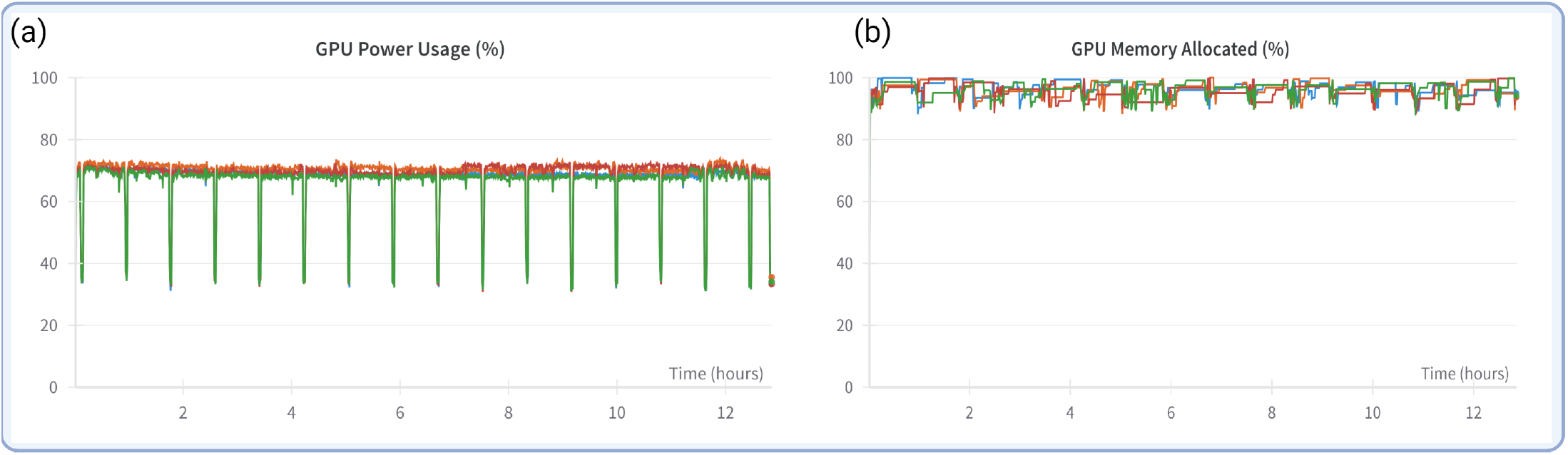
(a) Pretraining GPU Power Usage (a) GPU memory utilisation

**Fig. S3:**
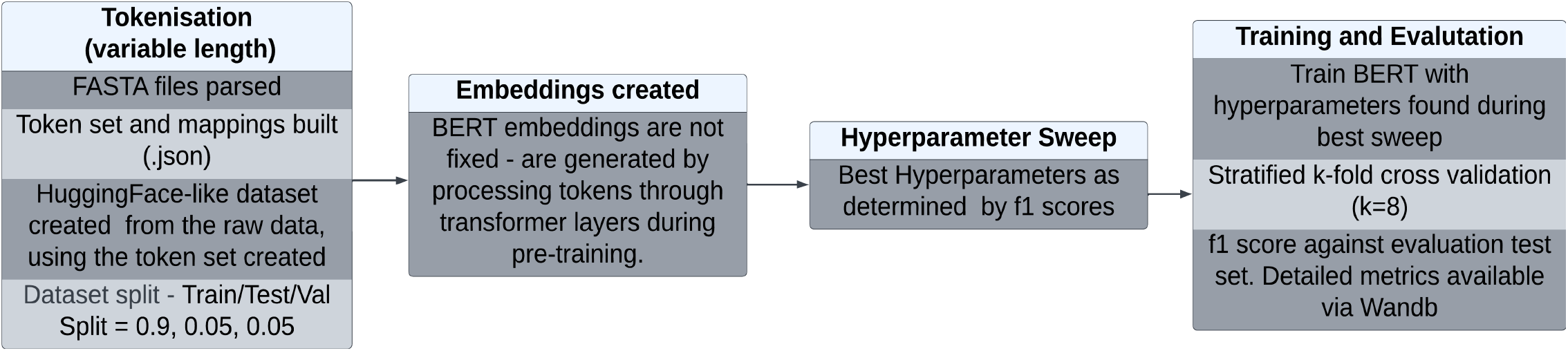
Empirical Token + BERT Workflow.

**Fig. S4:**
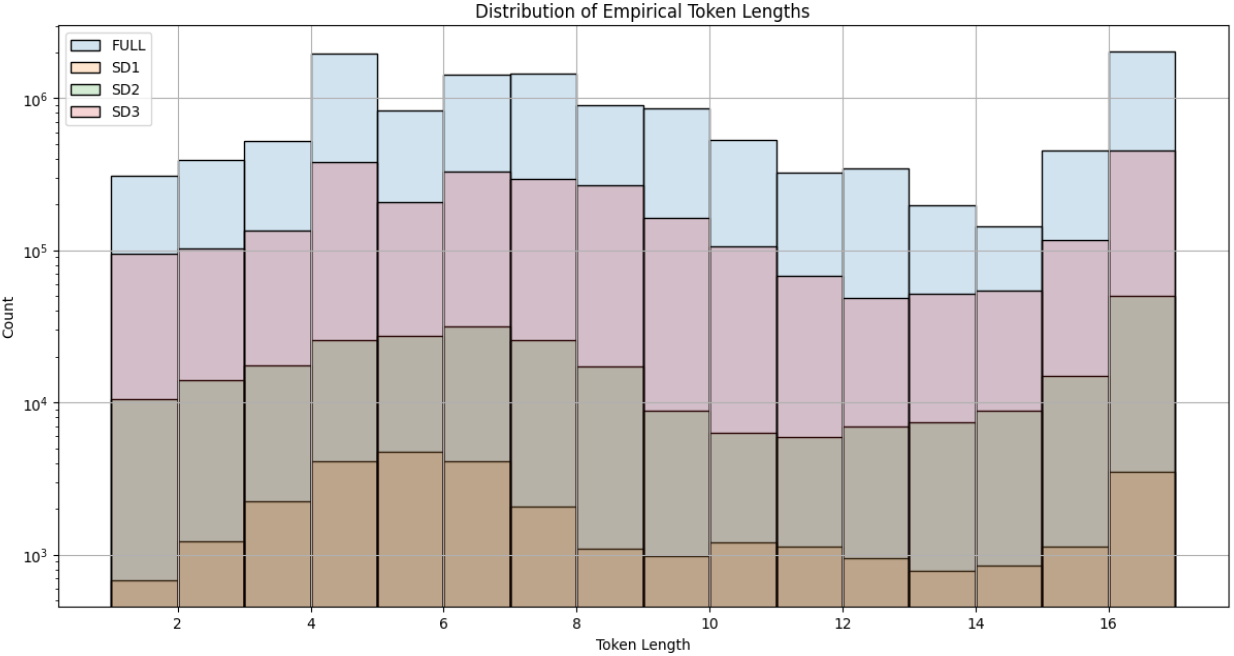
Distribution of Empirical Tokens - Length vs Count.

**Table. S1:**
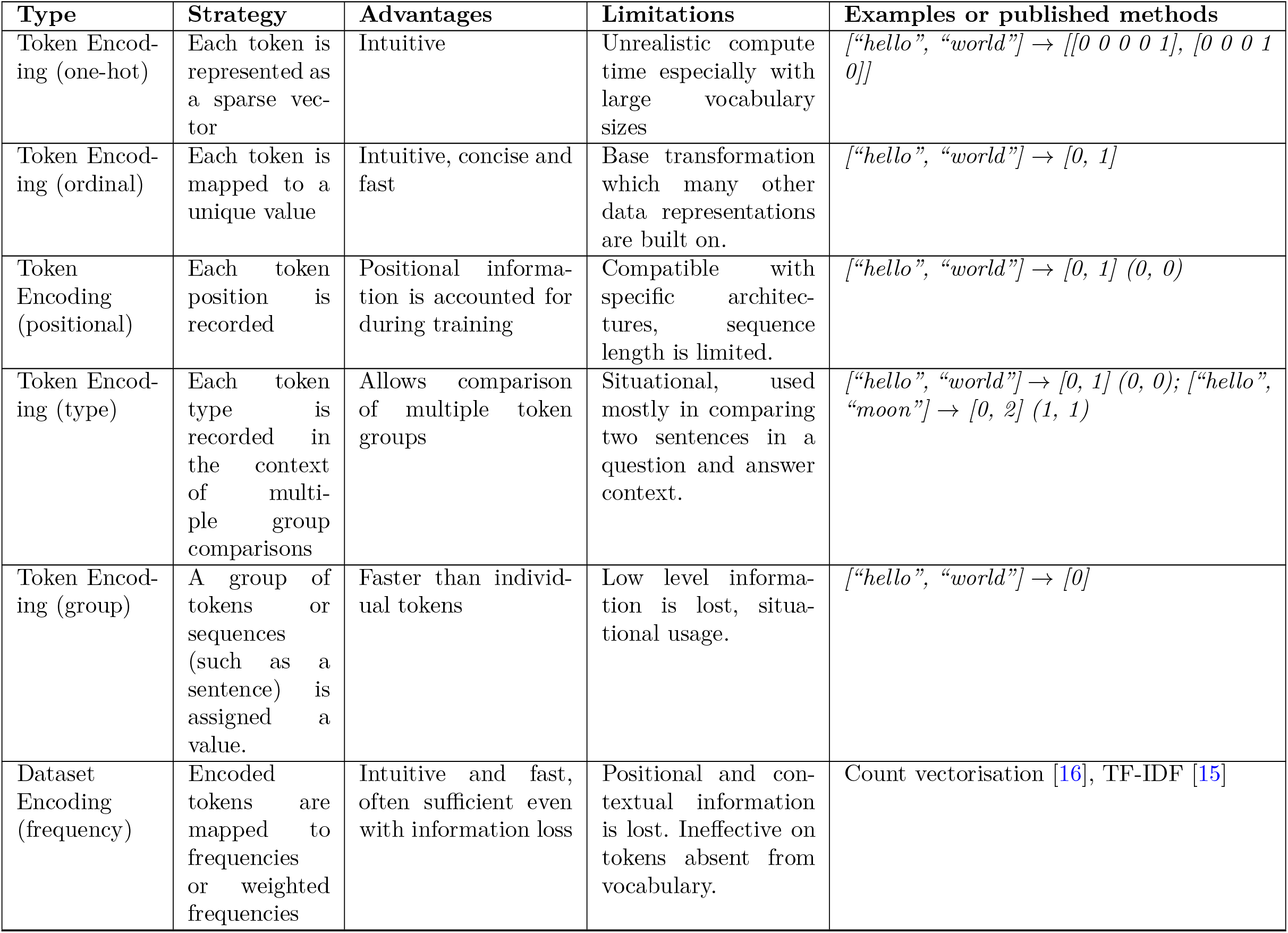
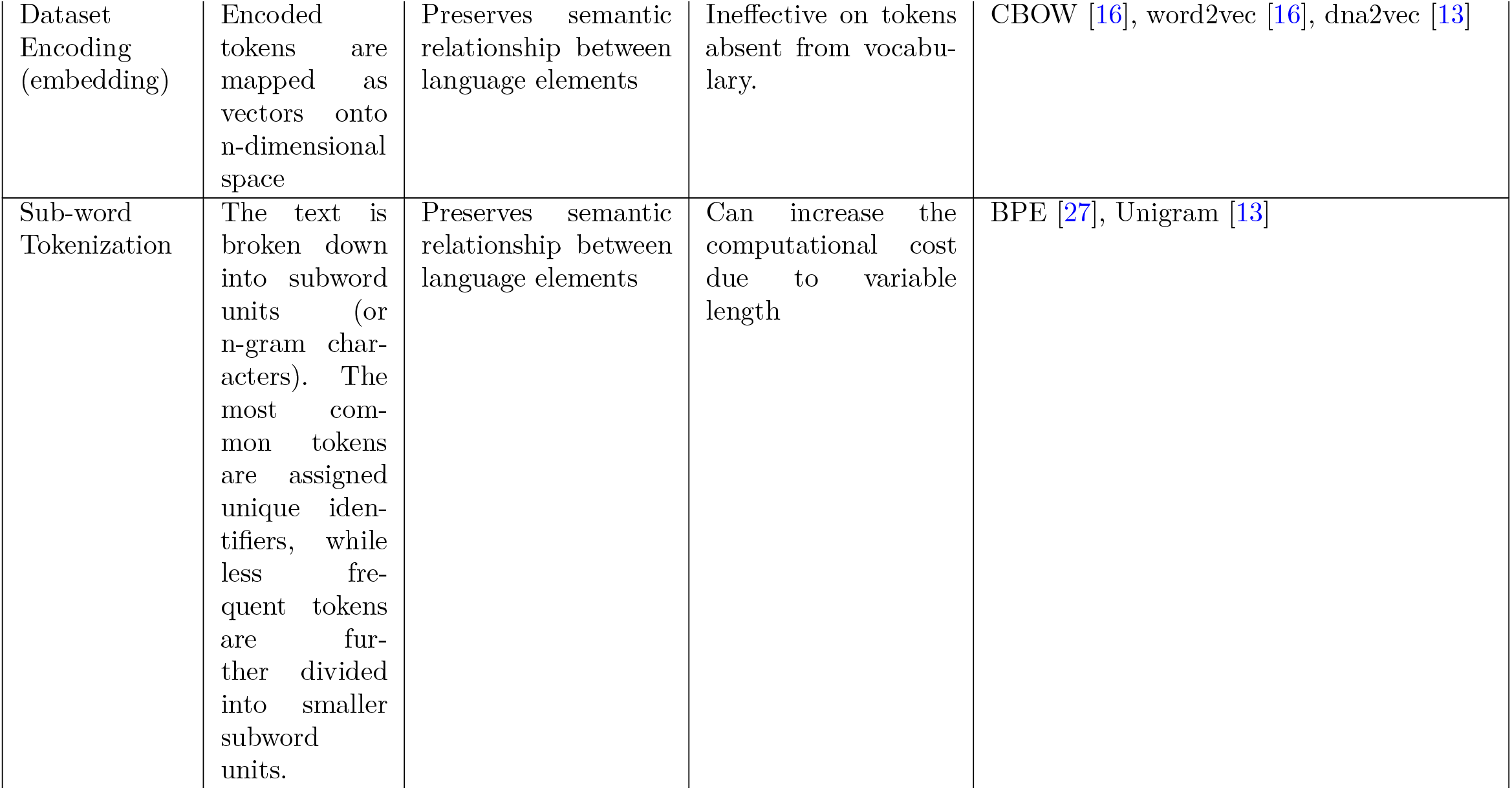
Comparison of Tokenisation approaches: Tokens are first recoded into a numeric form as a base. After this base step, more sophisticated transformations can be applied. Positional and group encodings are common in deep learning, while vector and frequency encodings are common in conventional machine learning.

**Table. S2:** Token counts: kmer9 vs Empirical.

| Data | Samples | Token count (Empirical) | K-mer Count(k=9) |
| --- | --- | --- | --- |
| SD1 | 1000/class | 30721 | 221271 |
| SD2 | 10000/class | 279225 | 2216539 |
| SD3 | 1,00,000/class | 2882443 | 22192432 |
| FULL | 4,50,000 | 12694670 | 98049771 |

