## Supplementary material for "*genomicBERT*: A Light-weight Foundation Model for Genome Analysis using Unigram Tokenization and Specialized DNA Vocabulary": Wandb

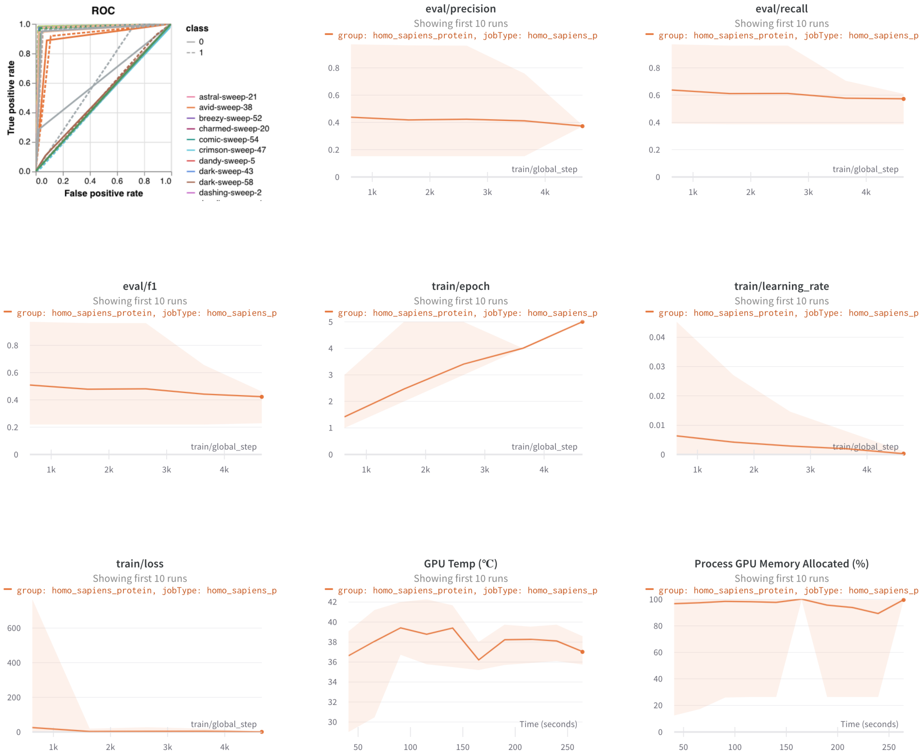


Figure S1: A sample of interactive plots generated and hosted on wandb showing various training, evaluation and compute metrics. This is not representative of the full range of plots that can be generated. New plots can be added as desired, and online versions of the plots are interactive.
